# Evaluation of the Luminex ARIES^®^ System for the Detection and Quantification of BK virus DNA in Plasma and Serum Samples from Kidney Transplant Recipients

**DOI:** 10.1101/344846

**Authors:** Tong Her, Ted E. Schutzbank

## Abstract

BKV nephropathy is as serious complication in renal transplant recipients due to the need for immunosuppression. Nearly 50% of renal transplant patients with BKV nephropathy experience a significant loss of function of the transplanted kidney. It is routine practice to screen renal transplant recipients regularly for BK viremia. In this study we compared the performance of BKV qPCR analyte specific reagents (ASR) by ELITech, and Luminex for measuring BKV viral load in plasma and serum using the Roche Cobas^®^ z480 instrument, and the Luminex ARIES^®^ platform respectively. A total of 34 patients previously tested on the z480, with results spanning the test ‘s linear range, were analyzed on the ARIES^®^. The BKV DNA copy number correlation between the two methods was very good with an R^2^ value of 0.98. The average difference in log copy number between the two methods was −0.3, indicating that the ARIES^®^ method may have slightly greater analytical sensitivity. BKV quantification results were closely matched between the two different methods. The workflow with the ARIES^®^ System is greatly simplified by elimination of DNA extraction and most hands-on steps. The high degree of automation allows samples to be tested as they arrive into the laboratory, resulting in enhanced patient care due to more rapid turnaround time for results back to the ordering physician.

## Introduction

BK virus (BKV), a member of the human polyomaviruses family, was first discovered in the urine of renal transplant patients with postoperative ureteral stenosis. (6) BKV is ubiquitous; in adults in the United States and Europe, seroprevalence ranges from 60% to 80%. Epidemiological studies indicate that BKV is contracted during childhood; between 3 to 4 years of age (11). Very little is known about the transmission of this virus or about events during primary infection. The majority of primary infections are asymptomatic; however, BKV may cause mild upper respiratory disease in young children (13). A latent infection is established in the kidneys of the infected host by BKV.

BKV is recognized as a significant cause of nephropathy in renal transplant patients and is a significant cause of graft failure in 1% to 10% of patients (7, 14). Approximately 40% of renal allograft recipients shed BKV in the urine, either transiently or continuously over weeks to months (1, 2). Nephropathy caused by BKV is asymptomatic and associated with increased serum creatinine levels. Nearly 50% of renal transplant patients diagnosed with BKV nephropathy experience a significant loss of function of the transplanted kidney. BKV viruria has also been demonstrated in approximately 10 – 25% of bone marrow transplant (BMT) patients, typically two months post-transplantation (2). The frequency of cystitis is higher in adult allogeneic compared to autologous BMT recipients. BKV can be demonstrated in the peripheral blood of both groups (5). Because detection of BKV in the urine of BMT patients in the absence of disease is common, it is difficult to correlate BKV as the cause of cystitis in these individuals (5).

Surveillance testing for BKV in the blood and urine of renal transplant recipients has become routine. In 2013, an expert panel recommended the use of either urine cytology or nucleic acid–based testing to screen renal transplant recipients monthly during the first 6 months post-transplantation, and then every 3 months until 2 years after transplantation (14). These guidelines included quantitative cutoffs for BKV loads that indicate additional testing. Urine DNA loads of >10^7^ copies/mL or plasma DNA loads of >10^4^ copies/mL that persist for more than three weeks constitute a diagnosis of “presumptive polyomavirus-associated nephropathy” and should be followed up with a renal biopsy (14).

There are no antiviral agents for the treatment of BKV. Treatment of kidney transplant recipients with BKV infection consists of lowering the dosage of immunosuppressive drugs used to prevent rejection of the transplanted organ to allow the patient’s immune response to clear the infection (3, 10, 12 16). For this reason, rapid turnaround time for quantitative BKV determination is important to initiate such treatment, and then to resume proper immunosuppressive therapy once the infection is cleared.

There are no FDA approved assays commercially available for BKV, so molecular testing is performed using laboratory developed tests; reagents for BKV nucleic acid amplification (NAAT) tests are available as either ASRs or RUOs from several commercial sources. In this paper we describe a rapid, sample-to-answer quantitative BKV quantitative polymerase chain reaction (qPCR) assay based on Luminex MultiCode^®^ PCR technology on the ARIES^®^ platform (Luminex Corp. Austin TX), which allows for samples to be analyzed as they are received in the laboratory as opposed to batch testing.

## Materials and Methods

### Clinical Samples

All clinical samples used in this study were obtained from kidney transplant recipients as part of routine screening for BKV viremia, submitted to the St. John Hospital and Medical Center Specialized Testing Laboratory for testing. All samples were de-identified prior to testing in this study eliminating the requirement for patient informed consent.

### Specimen processing

Plasma or serum samples from kidney transplant recipients were submitted to the St. John Hospital and Medical Center Specialized Testing Laboratory for BK qPCR virus testing. DNA was extracted from 1 mL of each sample using a bioMerieux NucliSENS^®^ easyMag^®^ nucleic acid extractor (bioMerieux, Durham, NC). The extracted nucleic acids from each sample were eluted into 50 µL of extraction buffer.

### BKV TaqMan qPCR

BKV qPCR testing based on TaqMan technology was performed using ELITech (ELITech Group, Logan, UT) 20X MGB Alert^®^ BK Virus Primers and 20X MGB Alert^®^ BK Virus Probe analyte specific reagents (ASR). The 25 µL reaction consisted of 1X of the primers, probe, MGB Alert^®^ Hot Start Master and MGB Alert^®^ BK Virus Internal Control primers and probes and 5 uL of the extracted nucleic acid solution. Analysis was performed using the Roche Cobas z480 platform with their proprietary user defined software package (Roche Diagnostics, Indianapolis, IN). Thermocycling conditions were 1 cycle each of 50°C for 2 minutes, 93°C for 2 minutes, followed by 45 cycles of 93°C for 15 seconds, 56°C for 30 seconds, and 72°C for 30 seconds. The standard curve was established using 5 µL of dilutions of a quantified synthetic BK virus DNA (ATCC, Manassas, VA) tested in triplicate.

### MultiCode^®^ PCR Analysis on the ARIES^®^ Platform

The MultiCode^®^ primer mix (Luminex Corp. Austin, TX) was prepared by adding 2 uL of MultiCode^®^ BK virus ASR primer and 2 µL of Control Primer (CP3) per sample to a 1.5 mL microcentrifuge tube. A volume of 4 µL of this primer mix was added to a MultiCode^®^ Ready Mix tube (Luminex, Austin, TX). The MultiCode^®^ Ready Mix tube contains all of reagents required for the MultiCode^®^ PCR reaction to occur. The prepared MultiCode^®^ Ready Mix tube is then attached to the appropriate position on the ARIES^®^ Extraction Cassette. Two hundred microliters (200 µL) of plasma or serum is added to the sample chamber of the cassette which is placed into the ARIES^®^ cassette holder and loaded into the ARIES^®^ instrument. Testing was performed according to standard instrument settings supplied by Luminex using their proprietary SYNCT™ software and the User Defined Protocol (UDP) application. To generate the standard curve, a 200 uL sample of each of the above BKV DNA dilutions was added to the sample chamber of the ARIES^®^ cassette. Each dilution was tested in duplicate.

## Results

### Generation of the BKV qPCR Standard Curve

The BKV qPCR standard curve on the Cobas z480 and ARIES^®^ platforms was generated using dilutions made from quantified synthetic BK virus DNA. The 6-member panel ranged from 1X 10^7^ to 5×10^2^ DNA copies/mL. Each dilution was tested in triplicate on the Cobas z480, and in duplicate on the ARIES^®^. The z480 software automatically generates a standard curve that can be saved and imported for subsequent analyses, meaning that a standard curve need not be generated for each run. The same dilution panel was used to generate the BKV DNA standard curve for the ARIES^®^ platform. In this case 200 uL of each panel member was added to ARIES^®^ Extraction Cassettes containing the MultiCode^®^ BK virus primers. The ARIES^®^ software was not designed for quantitative PCR analyses; it is necessary to import the standard curve Ct’s and patient data from the ARIES^®^ printout into Excel for analysis. The results, shown in Figure 2, panels A and B for the Cobas z480 and ARIES^®^ platforms respectfully are very similar. Based on the slopes of both standard curves the efficiency of amplification for the ELITech reagents on the z480, and the MultiCode^®^ BKV primers on the ARIES^®^ platform were determined to be very close at 97.12% and 99.18% respectively (where a slope of 3.22 = 100% efficiency (https://www.thermofisher.com/us/en/home/brands/thermo-scientific/molecular-biology/molecular-biology-learning-center/molecular-biology-resource-library/thermo-scientific-web-tools/qpcr-efficiency-calculator.html).

### Determination of the Limit of Detection (LoD)

The lower limits of detection (LoD) for both methods were determined by testing dilutions made from the quantified BKV DNA used to generate the standard curve. Each dilution was tested 20 times. The lowest dilution resulting in a positivity of ≥95% was set as the LoD. The LoD for the ELITech/z480 method was determined to be 500 copies/mL of original sample and 250 copies/mL for the MultiCode^®^/ARIES^®^ procedure (data not shown).

### Reproducibility of BK virus load testing on the ARIES^®^ platform

Five previously tested patient samples were used to determine reproducibility of the BK virus qPCR test on the ARIES^®^ platform. Three of these were positive and two were undetectable for BKV DNA. The results shown in Table 1 demonstrates a high degree of reproducibility as demonstrated by the very low percent coefficient of variation between each of the replicate analyses.

**Table 1:** Reproducibility of BKV qPCR on the ARIES^®^ Platform

| Sample | Replicate 1 |  | Replicate 2 |  | Replicate 3 |  | Avg. log cp/mL | SD | %CV |
| --- | --- | --- | --- | --- | --- | --- | --- | --- | --- |
|  | Ct | log cp/mL | Ct | log cp/mL | Ct | log cp/mL |  |  |  |
| 413531 | 32.2 | 3.99 | 31.6 | 4.17 | 31.0 | 4.35 | 4.17 | 0.18 | 4.32 |
| 412173 | *ND | ND | ND | ND | ND | ND | - | - | - |
| 410089 | 31.3 | 4.26 | 32.0 | 4.05 | 31.5 | 4.20 | 4.17 | 0.11 | 2.59 |
| 428857 | 26.2 | 5.79 | 25.8 | 5.91 | 25.9 | 5.88 | 5.86 | 0.06 | 1.07 |
| 341582 | ND | ND | ND | ND | ND | ND | - | - | - |

\*ND = Not detected

### Comparison of testing with clinical samples between the ELITech and ARIES^®^ qPCR assays

To compare the performance of both assays with actual clinical samples a panel of 34 plasma or serum samples submitted previously for BKV DNA qPCR testing were analyzed using the ELITech reagents on the z480 and the MultiCode^®^ BKV primers on the ARIES^®^ Platform. The results are shown in Table 1 and Figure 2. Test results for two different methods are considered equivalent if the log difference is ≤ 0.3 log copies/mL. The overall average Δ log difference between the ARIES^®^ and z480 BKV quantification tests was determined to be −0.3 logs. This value decreases to −0.24 if the samples with viral loads <500 copies per mL are removed from the analysis since 500 BKV DNA copies/mL is the lower limit of detection for the ELITech/z480 methodology. Linear regression analysis of the data in Table 1 (Figure 2 A) yielded a correlation coefficient (R-squared) value of 0.98, indicating very close agreement of the results obtained by both methods. The residual plot in panel B clearly demonstrates the validity of the linear regression plot in panel A.

## Discussion

Due to their highly immunocompromised state, kidney transplant recipients are at high risk of loss of their transplanted organ due to reactivated BK virus infection (1, 2, 7, 14). In the absence of antiviral agents that are active against BKV, “treatment” of BK virus infection in this population is limited to reduction of the use of immunosuppressive drugs to allow the patient’s immune system to clear the infection (10). As a result, the patient is placed at greatly increased risk of immunological rejection of the transplanted kidney (4, 9, 15, 17). Therefore, it is important that BKV viral load testing be performed as expeditiously as possible to determine the need for reduction of immunosuppression, with follow-up monitoring of BKV viral load to determine when appropriate levels of immunosuppressive therapy can be resumed (3, 8, 12, 16). Batch testing of plasma samples for BKV qPCR can result in a delay of testing, especially if the volume of testing is not sufficiently high to permit testing daily. Similarly, sending samples to reference laboratories can result in longer than desirable turn-around times to receiving results. In this study we evaluated the use of the Luminex MultiCode^®^ PCR primers on the ARIES^®^ real-time PCR platform to determine its applicability for performing quantitative PCR analysis, specifically for quantification of BKV DNA in plasma and serum samples. The ARIES^®^ BK virus test, utilizing primers based on MultiCode^®^ technology, was compared to our standard method with BKV PCR primers and an MGB probe from ELITech. For the latter, analysis was performed on the Roche Cobas z480 real-time PCR platform. Unlike the z480 platform, the ARIES^®^ instrument is not designed for qPCR analysis, so it is necessary to import the standard curve Ct’s and patient data from the ARIES^®^ printout into Excel for analysis. Since it would be neither convenient nor economically prudent to perform a standard curve each for each BK qPCR test run on the ARIES^®^, a standard curve was established using serial dilutions made from commercially sourced quantified BKV DNA, the results of which was stored and applied to successive test runs. The standard curves generated using both the ELITech and ARIES^®^ methods were nearly identical in terms of both slope and R-squared values (Figure 1). In addition, the slope of the standard curve generated on the ARIES^®^ platform of −3.34 demonstrates an amplification efficiency of >99% indicating a high degree of robustness of this assay design (19.). A new ARIES^®^ standard curve was generated when there was a lot change in any of the test reagents (ARIES^®^ test cartridges or MultiCode^®^ primers).

**Figure 1:**
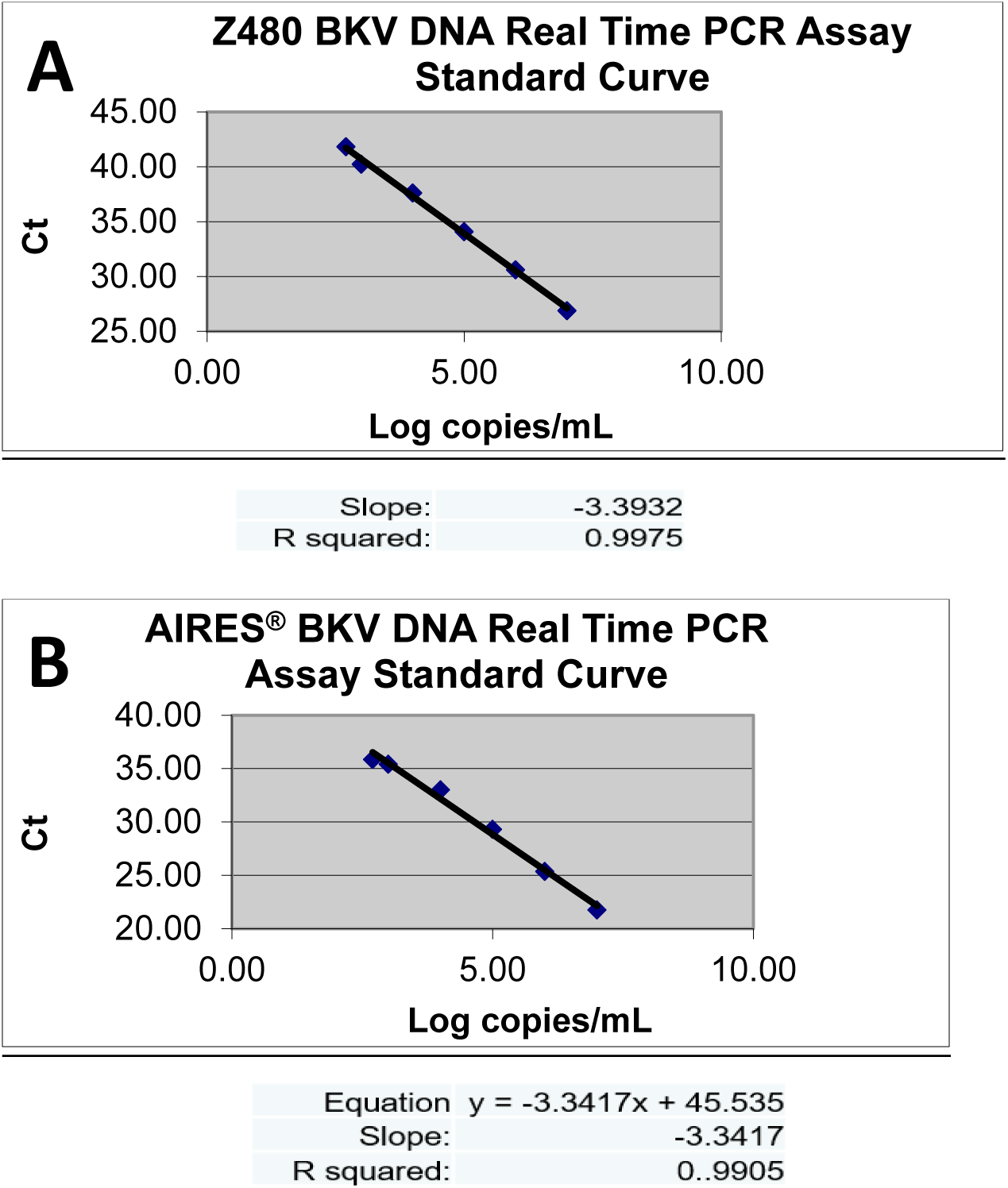
Generation of BKV qPCR standard curves Panel A – standard curve generated with serial dilutions of BKV purified DNA performed in triplicate. Panel B – standard curve generated with serial dilutions of BKV purified DNA performed in duplicate.

The robustness of this test design was clearly demonstrated by the results of the reproducibility experiment shown in Table 1. The log copy numbers of each of the samples in 3 different test runs resulted in low percent coefficients of variation demonstrating very low variability from run to run.

Comparison of testing with clinical samples between the ELITech and ARIES^®^ qPCR assays demonstrated essentially equivalent results between the two methods. The average difference between log copies per mL generated by both methods for each sample was −0.3. Linear regression analysis of the two sets of results (Figure 2) demonstrated 98% correlation between both methods. The high degree of correlation between results was somewhat surprising considering the large number of differences involved in the mechanics of performing both methodologies. The ELITech method on the z480 platform is entirely manual, including DNA extraction and assembly of the qPCR reactions. The ARIES^®^ method is completely automated, including the DNA extraction and reaction set-up processes. Another major difference is the qPCR chemistries used for both methods. The ELITech method utilizes TaqMan PCR technology and measures an increase in fluorescence with each PCR cycle. Due to its unique chemistry, MultiCode^®^ technology relies on loss of fluorescent signal per amplification cycle (18).

**Figure 2:**
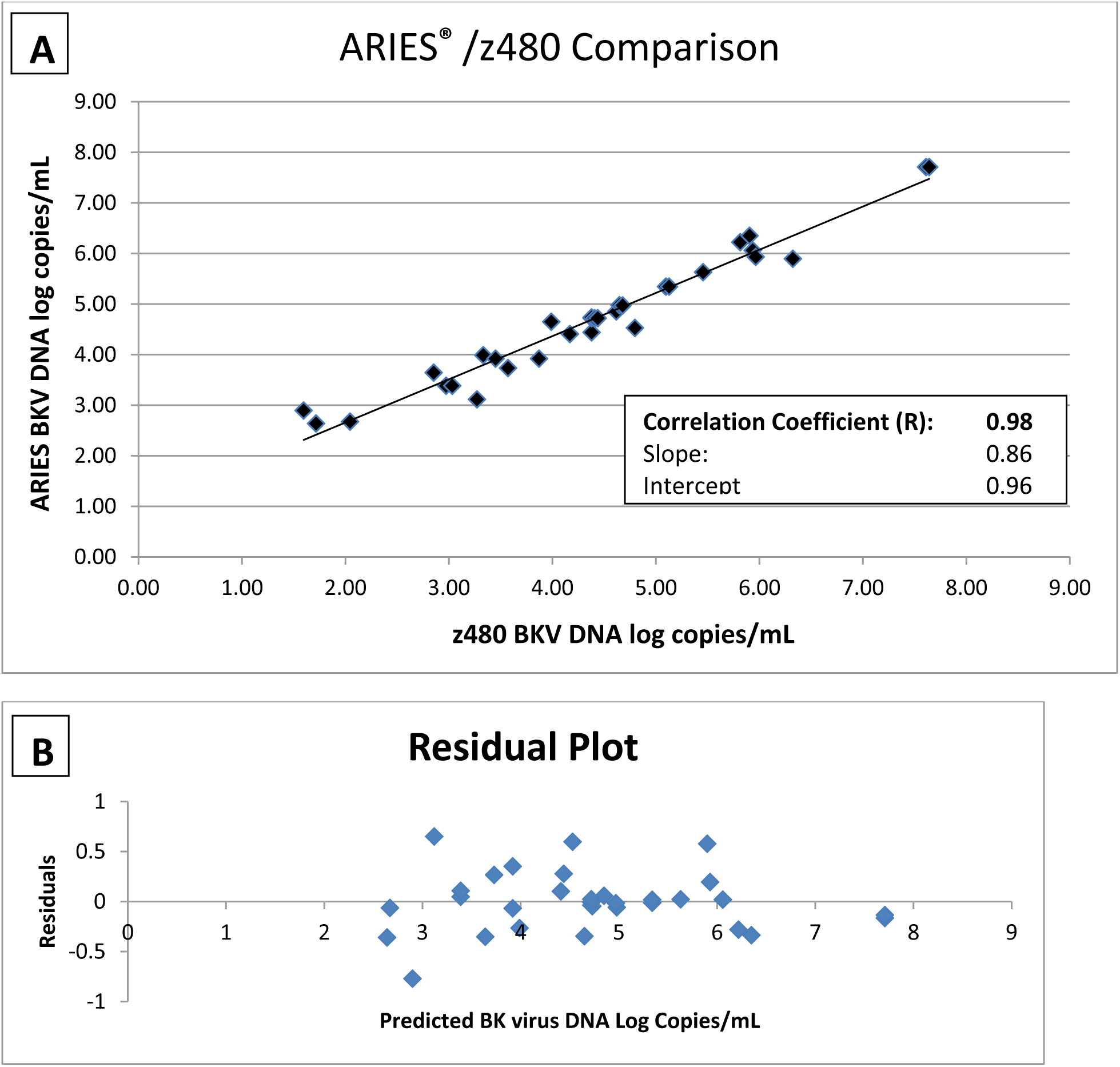
ARIES^®^ /z480 Comparison - Quantification of BK DNA in Clinical Samples Panel A – linear regression analysis of data from Table 2. Panel B – residual plot generated from regression analysis in panel A

**Table 2:** Comparison of testing with clinical samples between the ELI Tech and ARIES^®^ qPCR assays

| Sample | MultiCode®/ARIES | | ELITech/z480 | | $\Delta \log \text{LOG}$<br>Difference<br>ARIES®/z480 |
| --- | --- | --- | --- | --- | --- |
|  | Copies/mL | Log copies/mL | Copies/mL | Log copies/mL |  |
| 313611 | 39 | 1.6 | 787 | 2.9 | -1.3 |
| 318856 | 52 | 1.72 | 436 | 2.64 | -0.92 |
| 147329 | 79 | 1.9 | ND | ND | - |
| 416826 | 111 | 2.05 | 473 | 2.67 | -0.62 |
| 147329 | 254 | 2.4 | ND | ND | - |
| 497868 | 714 | 2.85 | 4,410 | 3.64 | -0.79 |
| 211229 | 1,079 | 3.03 | 2,427 | 3.39 | -0.36 |
| 299629 | 1,872 | 3.27 | 1,309 | 3.12 | 0.15 |
| 299521 | 2,149 | 3.33 | 9,756 | 3.99 | -0.66 |
| 162904 | 2,831 | 3.45 | 8,346 | 3.92 | -0.47 |
| 417959 | 3,729 | 3.57 | 5,406 | 3.73 | -0.16 |
| 162904 | 7,428 | 3.87 | 8,346 | 3.92 | -0.05 |
| 596591 | 9,785 | 3.99 | 44,607 | 4.65 | -0.66 |
| 308868 | 14,795 | 4.17 | 25,571 | 4.41 | -0.24 |
| 724744 | 23,966 | 4.38 | 27,553 | 4.44 | -0.06 |
| 531684 | 23,966 | 4.38 | 52,497 | 4.72 | -0.34 |
| 299531 | 23,966 | 4.38 | 54,309 | 4.73 | -0.35 |
| 234871 | 25,675 | 4.41 | 52,497 | 4.72 | -0.31 |
| 123044 | 41,590 | 4.62 | 71,244 | 4.85 | -0.23 |
| 596643 | 44,557 | 4.65 | 94,739 | 4.98 | -0.33 |
| 433255 | 47,735 | 4.68 | 93,462 | 4.97 | -0.29 |
| 40181 | 62,884 | 4.8 | 33,773 | 4.53 | 0.27 |
| 240660 | 125,252 | 5.1 | 221,263 | 5.34 | -0.24 |
| 355063 | 286,340 | 5.46 | 427,340 | 5.63 | -0.17 |
| 597242 | 654,604 | 5.82 | 1,671,599 | 6.22 | -0.4 |
| 473995 | 804,920 | 5.91 | 2,237,977 | 6.35 | -0.44 |
| 551855 | 862,338 | 5.94 | 1,150,917 | 6.06 | -0.12 |
| 40052 | 923,852 | 5.97 | 853,834 | 5.93 | 0.04 |
| 228172 | 2,112,027 | 6.32 | 787,062 | 5.9 | 0.42 |
| 149337 | 40,875,563 | 7.61 | 51,103,299 | 7.71 | -0.1 |
| 149337 | 43,791,366 | 7.64 | 51,103,299 | 7.71 | -0.07 |
| 433174 | ND | ND | 368 | 2.57 | - |
| 131845 | ND | ND | ND | ND | - |
| 293718 | ND | ND | ND | ND | - |
| Average $\Delta \log \text{copies/mL}$ | | | | | **-0.3** |

The high degree of correlation between both methods is important since it negates the need to re-baseline our kidney transplant population that have been previously tested with the ELITech methodology when switching to the ARIES^®^ BKV qPCR test.

In summary, our results clearly demonstrate that BKV quantification results were closely matched between the two different methods described above. The workflow with the ARIES^®^ System is greatly simplified due to elimination of DNA extraction and most hands-on steps. The high degree of automation allows samples to be tested as they arrive in the laboratory, resulting in enhanced patient care due to more rapid turnaround time for results back to the ordering physician.

